# Haplotype-resolved assemblies provide insights into genomic makeup of the oldest grapevine cultivar (Munage) in Xinjiang

**DOI:** 10.1101/2024.09.11.612401

**Authors:** Haixia Zhong, Xiaoya Shi, Fuchun Zhang, Xu Wang, Vivek Yadav, Xiaoming Zhou, Shuo Cao, Songlin Zhang, Chuan Zhang, Jiangxia Qiao, Zhongjie Liu, Yingchun Zhang, Yuting Liu, Hao Wang, Hui Xue, Mengyan Zhang, Tianhao Zhang, Yongfeng Zhou, Xinyu Wu, Hua Xiao

**Affiliations:** The State Key Laboratory of Genetic Improvement and Germplasm Innovation of Crop Resistance in Arid Desert Regions (Preparation), Key Laboratory of Genome Research and Genetic Improvement of Xinjiang Characteristic Fruits and Vegetables, Institute of Horticultural Crops, Xinjiang Academy of Agricultural Sciences, Urumqi, China; National Key Laboratory of Tropical Crop Breeding, Shenzhen Branch, Guangdong Laboratory of Lingnan Modern Agriculture, Key Laboratory of Synthetic Biology, Ministry of Agriculture and Rural Affairs, Agricultural Genomics Institute at Shenzhen, Chinese Academy of Agricultural Sciences, Shenzhen, China; College of Enology, Heyang Viti-Viniculture Station, Ningxia Helan Mountain’s East Foothill Wine Experiment and Demonstration Station, Northwest A&F University, Yangling 712100, China

**Keywords:** viticulture, genome assembly, somatic mutation, Munage grape, anthocyanin

## Abstract

Munage, an ancient grape variety that has been cultivated for thousands of years in Xinjiang, China, is recognized for its exceptional fruit traits. There are two main types of Munage: white fruit (WM) and red fruit (RM). However, the lack of a high-quality genomic resources has impeded effective breeding and restricted the potential for expanding these varieties to other growing regions. In this study, we assembled haplotype-resolved genome assemblies for WM and RM, alongside integrated whole genome resequencing (WGS) data and transcriptome data to illuminate specific mutations and associated genes in Munake and the genes associated with fruit color traits. Selective analysis between Munage clones and Eurasian grapes suggested that adaptive selection exists in Munage grapes, with genes enriched in processes including cell maturation, plant epidermal cell differentiation, and root epidermal cell differentiation. The study examined the mutations within Munage grapes and found that the genes *PMAT2* on chromosome 12 and *MYB123* on chromosome 13 are likely responsible for color variation in RM. These findings provide crucial genetic resources for investigating the genetics of the ancient Chinese grape variety, Munage, and will facilitate the genetic improvement in grapevine.

## Introduction

The cultivated grapevine (*Vitis vinifera* L.) is closely associated with human civilization. The genetic diversity of grapevine cultivars has allowed this fruit crop to be utilized for be used for various purposes, and this plant has also become emblematic of cultural identity in many regions due to its central role in food and social practices. (Banilas et al., 2009). The Eurasian grape was first domesticated and cultivated at the region spanning from the eastern coast of the Mediterranean Sea to the Caucasus, which lies between the Caspian Sea and the Black Sea. (Myles et al. 2011; Zhou et al. 2017a; Xiao et al., 2023; Dong et al., 2023), and then introduced to Europe, Asia, and other regions (Maghradze et al., 2020). It appeared in Central Asia around 3000 B.C. and was introduced in Xinjiang, China, via Iran in around 1000 B.C. (3000 years ago) (Zhao et al., 2019). There are no naturally occurring Eurasian species of grapes in China, and Chinese Eurasian species of grapes were introduced from Central Asia to Xinjiang, via the early Silk Road, from which they were gradually introduced in other regions in China. Archaeological discoveries showed that grape seeds from Central Asia reached Xinjiang 2500 years ago (Li, 2021).

Xinjiang has a long history of grape cultivation and has inherited a rich variety of ancient native grape germplasm resources with irreplaceable genes due to its long history of cultivation (Li, 2015; Zhang et al., 2022). The area covered by old native varieties such as Thompson Seedless, Mare Teat (also known as Milk), and Munage grapes has reached 80,000 hectares, which is relatively stable, occupying about 10% of the total area. A thousand-year cultivation history signifies the unique characteristics of these varieties. Over the course of more than 2500 years of extensive selection and cultivation, many variations have accumulated, including both beneficial and detrimental ones. These variations in germplasm resources are important for breeding or screening of new varieties. However, the trait-specific genes of these varieties have not yet been researched and utilized. Munage grapes are the most characteristic and high-quality native varieties in Xinjiang (Liu et al., 2023). These grapes are sought after for their large fruits, delicate skin, and juicy flesh, making them a popular variety of table grapes. The grapes are further categorized into two types based on the color of their skin: red Munage (RM) and white Munage (WM). The ancient variety Munage has unique regional adaptability, and its excellent traits and specific genetic resources need to be excavated and utilized urgently. Because of its excellent genetic diversity, it is still the main variety in the Xinjiang region. In addition, it is assumed that RM grapes are bud mutant of WM, but lack of clear scientific evidence presently. Bud sprout mutation is a common phenomenon in fruit plants and plays a vital role in improving the overall quality of grapevines. For instance, a famous Chinese variety Jinzao Wuhe is a bud sprout of Himord Seedless with enlarged barriers (Huang et al., 2023). Moreover, Thompson seedless is also assumed to be a shoot mutant of Sultanina, which was further distributed and maintained by cuttings (Ledbetter and Ramming, 1989). The origin and domestication history of Munage remain unclear. The color variation observed in Munage presents a significant opportunity for us to utilize this genetic resource to gain a better understanding of the complex mechanisms involved in berry color development.

The fruit color is among the most important factors affecting the acceptability of grapes. The red pigment of grapes basically comes from the anthocyanins, which not only determines the color and commerciality of grapes but, as important flavonoid secondary metabolites, also improves stress resistance. However, the mechanism of grapevine fruit anthocyanin synthesis is still not very clear. The available evidence shows that anthocyanins are downstream products of the phenylpropanoid pathway. The studies related to color development in grapes showed that grapevine berry anthocyanin biosynthesis is mainly dependent on multiple factors, including environment, water, phytohormone, etc. (Li et al., 2024; Teixeira et al., 2013). The available information about the role of molecular mechanisms reveals that chalcone synthase (CHS), dihydroflavonol 4-reductase (DFR), myeloblastosis protein (MYB), basic helix-loop-helix (bHLH), WD, as well as MADS-box and SQUAMOSA promoter-binding protein (SBP-box) genes regulate berry color in grapes and other plants (Jaakola, 2013). Multiple studies have substantiated the important role of MYB genes in berry color development. These studies have established that MYB transcription factors have a regulatory influence on berry color (Koyama et al., 2014; Xie et al., 2020). The MybA1 and MybA2 genes have been identified as regulators of color development in some grape varieties (Jiu et al., 2021). *MybA1* is primarily expressed in the grape pericarp tissue, where it plays a crucial role in accumulation of anthocyanins (De Lorenzis et al., 2016) whereas, *MybA2* has been demonstrated to significantly activate the UFGT promoter, leading to a remarkable increase in expression level (Bogs et al., 2007). Moreover, the *MybA* transcription factor gene family is believed to play a crucial role in determining the range of anthocyanin concentrations found in grape skins. Further research has clarified the twofold role of bHLH transcription factors. First, they directly regulate the expression of DFR. Secondly, they interact with MYB, promoting the increased accumulation of color compounds (Walker et al., 2007). Some recent studies have substantiated the role of WD40 transcription factors in the regulation of anthocyanins in grape berries (Li et al., 2021). Moreover, the expression levels of flavanone 3-hydroxylase (F3’H) and flavonoid 3’,5’-hydroxylase (F3’5’H) determine the ratio of different anthocyanins, which in turn affects skin color. Also, decreasing the expression of FLS or genes related to the production of proanthocyanidins in grapevine berries can increase the accumulation of anthocyanidins (Azuma et al., 2012). However, there is a lack of studies that explore the detailed mechanisms about the complex genetic network involved in color development.

The progress of high-throughput sequencing technologies has greatly accelerated the identification of genetic variations, including small insertions/deletions (InDels) and single nucleotide polymorphisms (SNPs). More recently, genome sequencing in grapes has greatly contributed to our understanding of the structure of the *Vitis* genome (Wang et al., 2024). In the past decade, the utility of haplotype assemblies in identifying genes associated with agronomic traits has been well demonstrated in grapevine (Shirasawa et al., 2022; Cochetel et al. 2023; Zhang et al., 2023; Calderón et al., 2024; Wang *et al*., 2024; Zhang et al. 2024). In this study, we generated high-quality haplotype-resolved genome assemblies of the ancient Chinese grape variety Munage. Comparative genomics was employed to identify selected differential segments and structural variations (SVs) in the genomes of different varietal types of the ancient grape variety Munage grape. This allowed us to identify the potential functions of genes, elucidate the evolutionary relationships, and explore the structural variations of the genomes of ancient grape varieties from Xinjiang, China. Additionally, we examined the expression patterns of the transcriptome in various Munage grape tissues to identify the superior traits and specific genes associated with these grapes, particularly those related to color. By mining key color genes for fruit quality traits, we aim to provide genetic resources and a molecular foundation for enhancing ancient grape varieties. This will add in the accurate identification of essential agronomic traits and facilitate the genetic selection of superior ancient grape varieties. Moreover, it will offer genetic resources for future grape genome designs, serving as a scientific basis for developing new grape germplasm and advancing breeding technology.

## Results

### Haplotype-resolved assemblies of Munage genome

The plant materials used in this study were from an 80-year-old clonally propagated individual of white Munage (WM) from Xinjiang, China, and an individual of red Munage (RM) with a mother tree over 30 years old (Fig. 1A). To gain a deeper understanding of WM and RM, we performed PacBio sequencing on both RM and WM, obtaining 64.27 Gb of HiFi reads for WM and 49.78 Gb of HiFi reads for RM. Hi-C data were sequenced using the Illumina HiSeq X Ten platform, resulting in 52.35 Gb of Hi-C reads for WM and 52.67 Gb of Hi-C reads for RM.

**Fig. 1.**
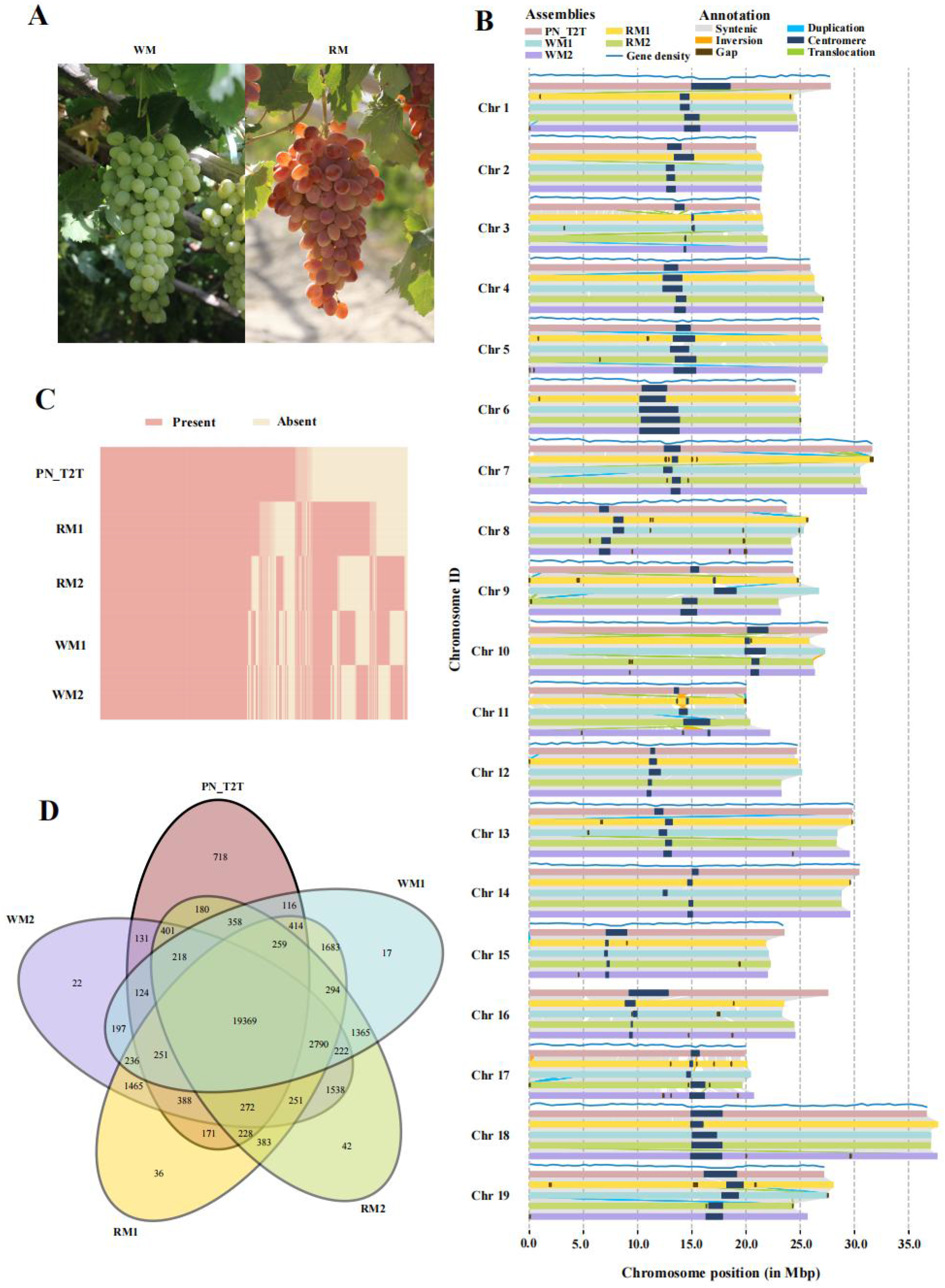
(A) Collinearity alignments between the PN_T2T genome and the WM1, WM2, RM1 and RM2 genomes. (B) Fruit images of RM and WM. (C) Visualization of the distribution of homologous genes in the PN_T2T, WM1, WM2, RM1, and RM2 genomes. (D) Differences in homologous genes among the PN_T2T, WM1, WM2, RM1, and RM2 genomes.

We initially estimated the genome sizes of the two species using the k-mer metric method with HiFi reads. The results indicated that the genome size of WM is approximately 537.74 Mb, while the genome size of RM is approximately 461.60 Mb (Fig. S1A, Table S1). We performed an initial assembly using HiFiasm, which yielded four haplotypes for the two species at the contig level. The contig N50 values were 24.32 Mb for haplotype 1 of WM (WM1), 26.33 Mb for haplotype 2 of WM (WM2), 15.55 Mb for haplotype 1 of RM (RM1), and 17.08 Mb for haplotype 2 of RM (RM2) (Table S2).

Through Hi-C data integration, we obtained haplotype-resolved sequence assemblies for white and red Munage and performed gene annotation and transposable element (TE) annotation. The sizes of the assemblies were 488,103,118 bp for WM1, 48,935,377 bp for WM2, 489,516,681 bp for RM1, and 480,619,624 bp for RM2. We annotated 33,942, 34,034, 35,292, and 35,007 genes for WM1, WM2, RM1, and RM2, respectively, with TE annotations of 67.04%, 67.15%, 67.02%, and 66.56%. The BUSCO completeness scores were 98.3%, 98.6%, 98.3%, and 98.5%, respectively (Fig. S1B).

Our assembled haplotypes were compared with PN_T2T (Shi et al., 2023). The results showed high collinearity among the five haplotypes (Fig. 1B). However, there were also noticeable differences among the haplotypes. By comparing the gene annotations among the five haplotypes, 718 orthologous genes unique to PN_T2T were identified. These genes were mainly enriched in pathways related to secondary metabolism, defense response, asymmetric cell division, and secondary metabolite biosynthesis during embryonic development (Fig. 1C, Table S4). Further, we identified 22 unique orthologous genes in WM1, which were mainly enriched in pathways such as response to stimulus, response to external biotic stimulus, response to biotic stimulus, biological processes involved in interspecies interactions, and obsolete multi-organism processes (Table S5). RM1 was identified to have 36 unique orthologous genes, primarily enriched in pathways like obsolete cellular nitrogen compound metabolic processes, obsolete RNA polyadenylation, obsolete nitrogen compound metabolic processes, RNA 3’-end processing, and RNA metabolic processes (Table S6). RM2 has 42 unique orthologous genes, which are mainly enriched in pathways such as plant organ development, RNA splicing via transesterification reactions with bulged adenosine as a nucleophile, RNA splicing via transesterification reactions, and RNA splicing (Fig. 1D, Fig. S2A-D, Table S7).

### Population Subdivision

To confirm the genetic history of Munage, we selected and analyzed resequencing data from 59 grape samples. The samples included 10 wild grapes (*V. vinifera* ssp. *sylvestris*) from Europe (EU), 10 wild grapes from the Middle East and Caucasus region (ME), 20 domesticated grapes (*V. vinifera* ssp. *vinifera*), and 9 Munage grapes from Xinjiang (Table S3). Additionally, three muscadine grapes were used as an outgroup.

After calling and filtering SNPs using PN_T2T as a reference genome, we constructed a phylogenetic tree of the samples using the general time-reversible (GTR) model in FastTree software (Price et al., 2010). In the resulting phylogenetic tree, it is evident that wild grapes and domesticated grapes form two distinct branches, indicating early divergence between them (Fig. 2A). The domesticated grapes, including Eurasian grapes (table, grapes for fresh consumption, and wine, the grapes for wine production) and Munage, show a clear monophyletic structure, indicating a common origin. Principal Component Analysis (PCA) analysis also confirmed the differentiation among these populations (Fig. 2B). Notably, high value (0.98) of identity-by-descent (IBD) analysis between WM and RM revealed that one of the cultivars originated by the bud mutation of the other (Fig. S3). We marked the distribution points of Munage in Xinjiang on the map and combined it with climate conditions to simulate the distribution trend map of Munage (Fig 2C). According to the simulation results, the distribution of Munage gradually decreases from the west to the east of Xinjiang. The central part of Xinjiang has a lower count, mainly due to the presence of the Taklamakan Desert, which is unsuitable for plant growth (Fig. 2C).

**Fig. 2.**
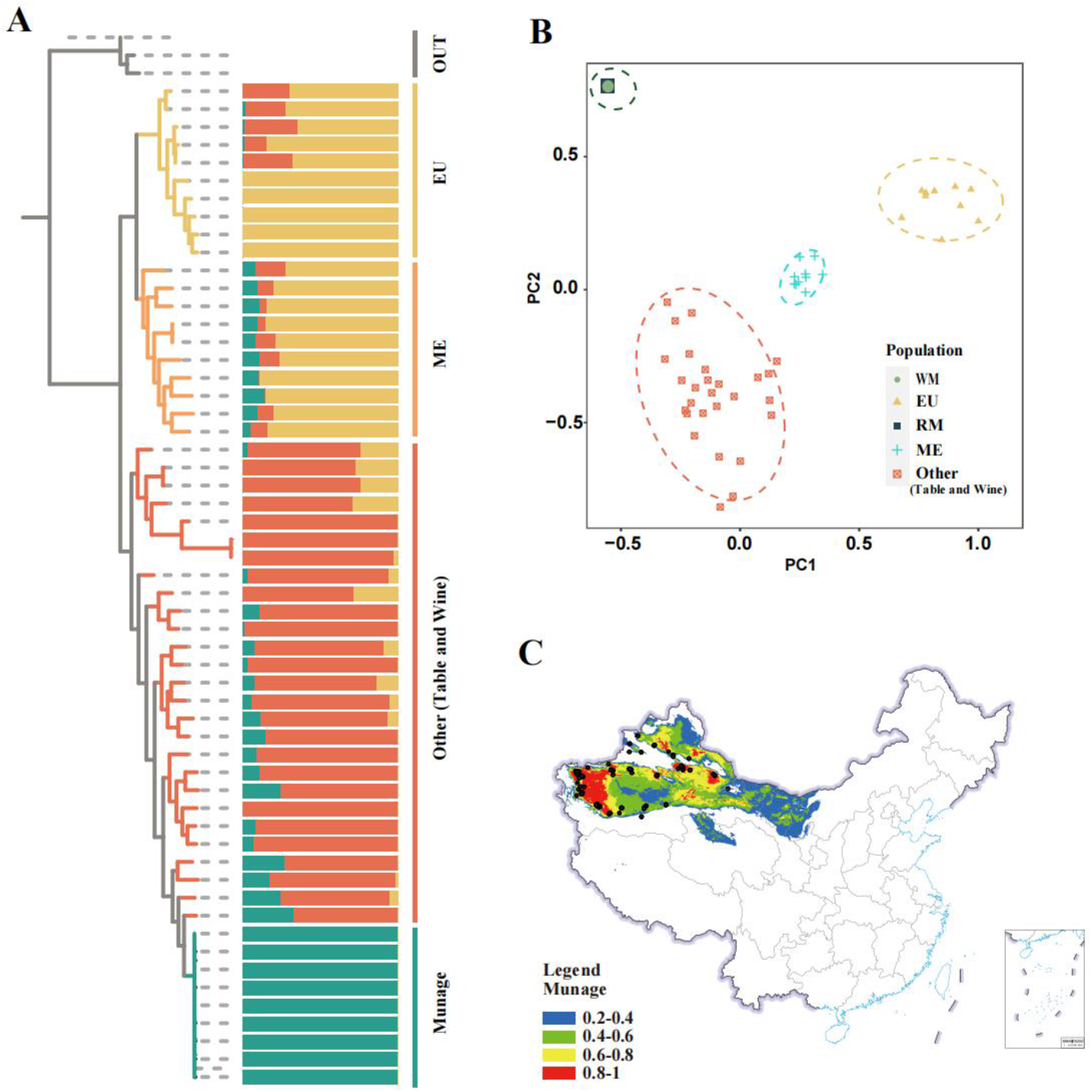
(A) Phylogenetic tree with admixture analysis, where branches represent different groups. The admixture plot shows K = 4. (B) PCA analysis based on SNP loci from 56 re-sequenced samples. (C) Predicted distribution of Munage based on climate, with black dots indicating sampling points.

### 3.3 Selective sweep in the Munage grape

We identified the genomic regions that underwent selective sweeps to study the domestication lineage between Munage and other domesticated grape populations. In Population Branching Structure (PBS) analysis, we designated Munage as Population One, other domesticated grapes as Population Two, and ME as Population Three. This analysis was carried out to reveal the differences between Munage and other wine and table grape populations in specific genes or genomic regions, and to identify the regions that have been influenced by selection during the ongoing domestication process. The PBS results showed significant differentiation in many regions across 19 chromosomes (Fig. 3A). We selected the top 5% of regions with the highest PBS values for Gene Ontology (GO) enrichment analysis, finding that these regions are mainly enriched in pathways such as cell maturation, plant epidermal cell differentiation, root epidermal cell differentiation, root hair cell development, glucan biosynthetic process, trichoblast differentiation, root hair cell differentiation, trichoblast maturation, cellular metabolic processes, and compound salvage (Fig. 3C, Table S8). These pathways are closely related to key physiological processes and the maintenance of vital activities in plants, which are likely associated with adaptation to the specific environment in Xinjiang.

**Fig. 3.**
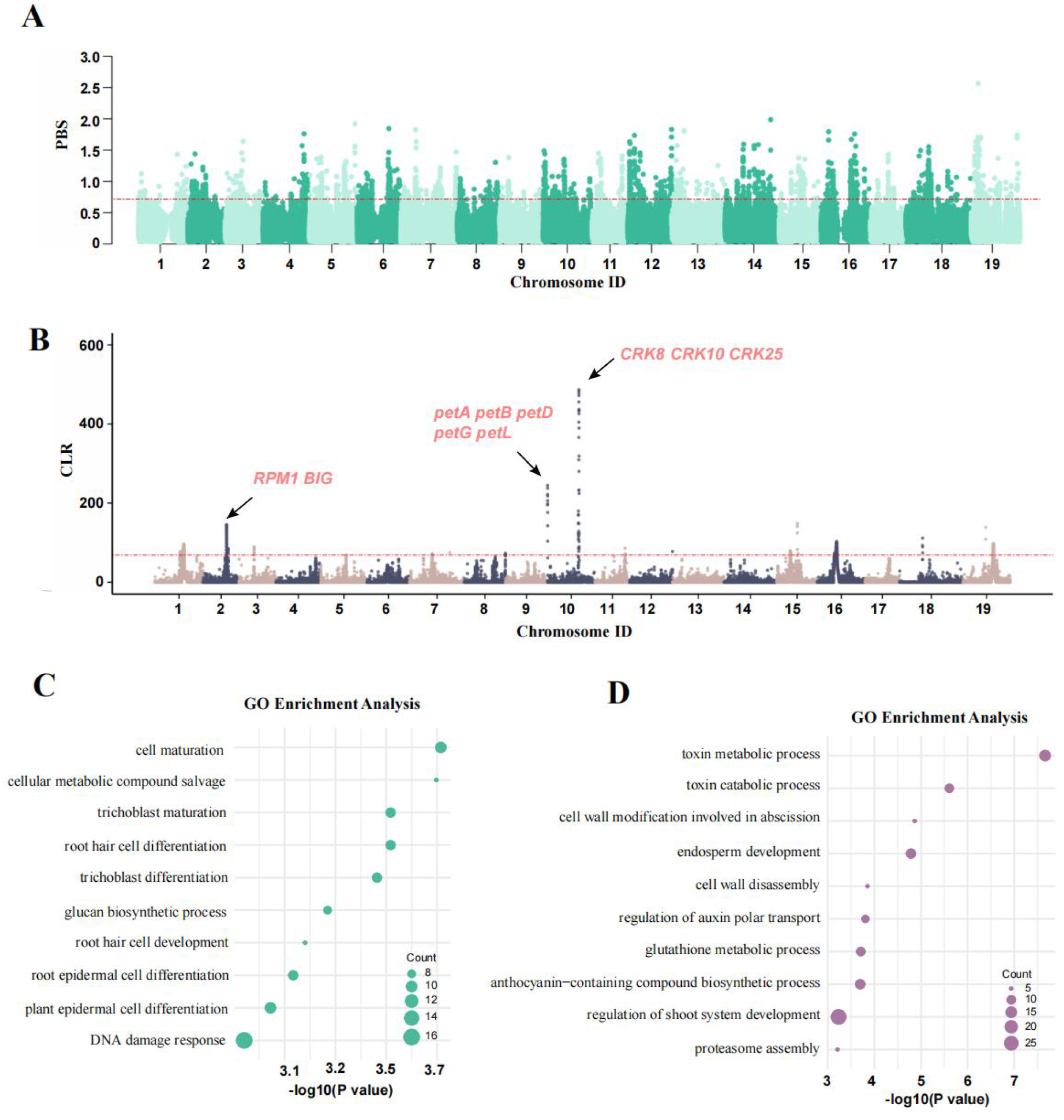
(A) PBS analysis based on SNP variant information, using the MUNAGE, Other, and ME groups. (B) SweeD analyses of the MUNAGE group. (C) GO term enrichment of the biological process in the top 1% of genes in the PBS analyses. (D) GO term enrichment of the biological process in the top 1% of genes in the SweeD analyses.

To further explore the selection signals within the Munage population during domestication, we performed composite likelihood ratio (CLR) analysis on nine Munage grapevine samples using SweeD software. Through the CLR analysis of the Munage population, we detected several selected genomic regions, particularly on chromosomes 2 and 10 (Fig. 3B). On chromosome 2, the *RPM1* gene (Boyes et al., 1998) was located at the selection region. *RPM1* is a classical plant resistance gene (R gene), encoding a protein that recognizes pathogen effectors and activates the plant immune system to resist pathogen invasion. Another identified gene is the *BIG* gene (Gil et al., 2001), which is involved in the regulation of plant hormone signaling, such as auxin and gibberellin. It modulates cell sensitivity and response to these hormones, affecting plant growth and development. On chromosome 10, a gene cluster including *petA, petB, petD, petG*, and *petL* (Cramer, 2019) was identified, all of which encode proteins related to the cytochrome b6f complex in the photosynthetic electron transport chain, playing crucial roles in photosynthesis. Additionally, *CRK8, CRK1*, and *CRK25* genes (Yadeta et al., 2017) were detected, which are cysteine-rich receptor-like kinases (CRKs) that play specific roles in plant immune responses to various pathogens and other stresses. We also conducted GO enrichment analysis on the top 1% regions with the highest composite likelihood ratios, which were primarily enriched in processes such as camalexin metabolic process, regulation of shoot system development, anthocyanin-containing compound biosynthetic process, glutathione metabolic process, regulation of auxin polar transport, cell wall disassembly, endosperm development, cell wall modification involved in abscission, toxin catabolic process, and proteasome assembly (Fig. 3D, Table S9). These pathways are closely related to the regulation of normal physiological activities in plants.

### Effects of somatic mutations on color change

It is commonly believed that RM arose as a bud mutant from WM, which also been demonstrated in this study above. WM has green shoots with a slight brownish tint, crispy and sweet fruit flesh, while RM has green shoots with a slight reddish-purple color, reddish branches, and crispy, slightly tart fruit flesh. To further investigate the differential effects of the bud mutation on WM and RM, we first used SNP variation (Fig. 4A) data from six RM samples for sequence similarity (*D*_*xy*_) analysis, measuring the absolute difference in nucleotide diversity. We then performed GO enrichment analysis on the genes in the top 5% of regions based on *D*_*xy*_ values and found that they were mainly concentrated in pathways such as RNA modification, response to salt stress, response to water deprivation, and cell death (Fig. S4A). Additionally, based on gene functional annotation, we identified a homologous gene of *Nicotiana tabacum, PMAT2*, on chromosome 12, which is related to anthocyanin synthesis (Fujiwara et al., 1997). Previous research suggests that this gene is involved in anthocyanin synthesis, which imparts the color of flowers, fruits, and leaves. Furthermore, we have identified the *MYB123* gene on chromosome 13, which controls the synthesis of anthocyanins in specific tissues of the inner pericarp. This gene may influence the skin color of Munage fruits (Wang et al., 2019b). Concurrently, we integrated RNA-seq data from RM and WM peels for temporal analysis of differentially expressed genes. A total of 578 differentially expressed genes were identified, including 216 up-regulated genes and 362 down-regulated genes (Fig. 4B). Upon analyzing the up- and down-regulated genes, we found that the up-regulated genes were primarily enriched in pathways related to water transport, fluid transport, transmembrane transport, and solute transport (Fig. S4B, Table S10). Additionally, pathways specific to skin color, including pigment metabolism and biosynthesis pathways, were also identified. In contrast, the down-regulated genes were enriched in pathways related to heat response, temperature stimulus response, pigment metabolism, and pigment biosynthesis processes (Fig. 4C, Table S11). This indicates that the expression of these genes decreases in WM, resulting in the green coloration of the fruit peel.

**Fig. 4.**
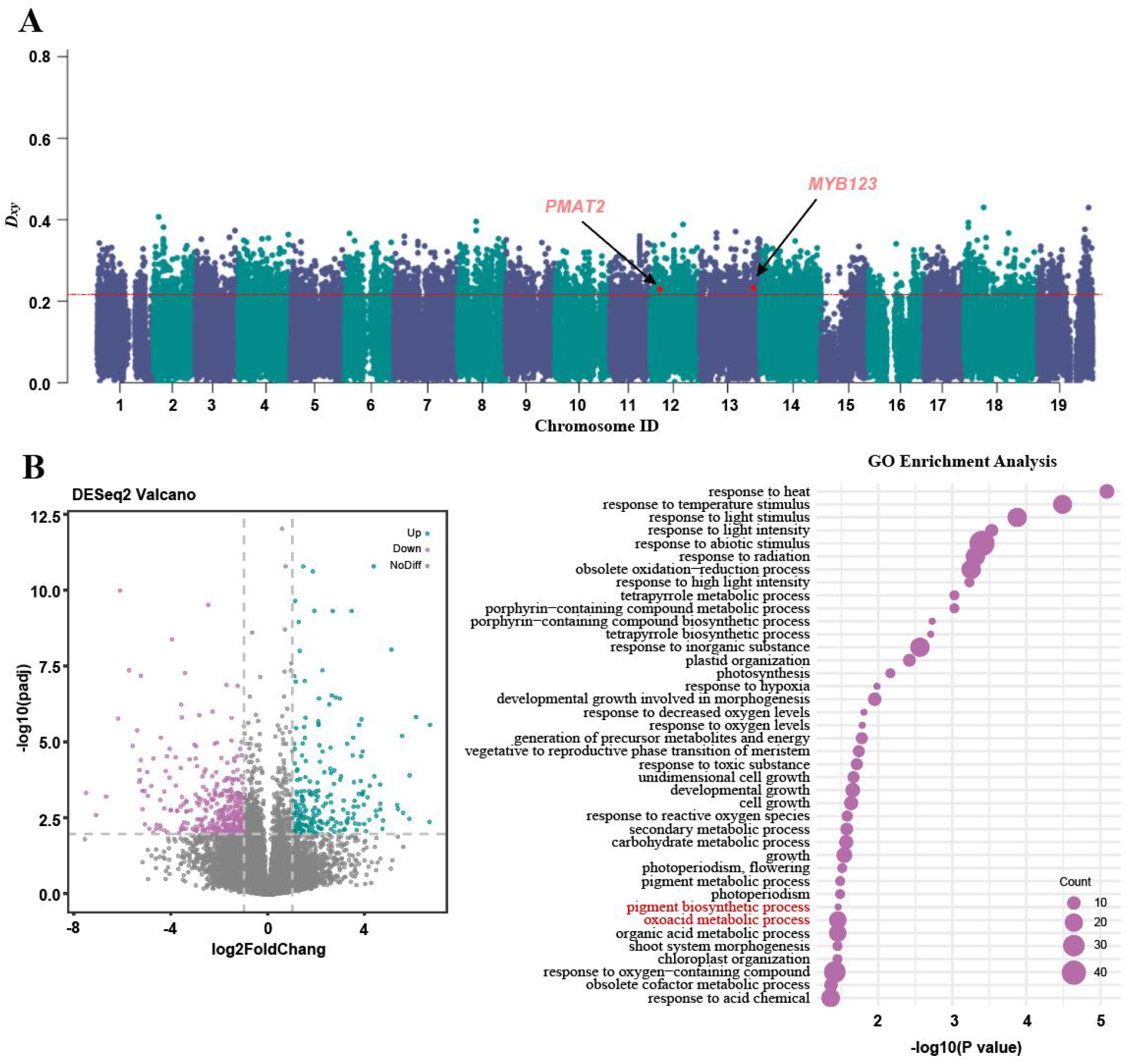
(A) *D*_*xy*_ analysis of the RM group. (B) Differential gene expression analysis based on WM and RM transcriptome data. (C) GO enrichment analysis based on downregulated differentially expressed genes.

## Discussion

Genomes are essential for research and breeding as they provide comprehensive insights into genetic variations, enabling the development of improved traits and optimized crop varieties. In 2024, the first telomere-to-telomere grape genome assembly was published using PN40024 (Shi *et al*., 2023), a highly homozygous cultivar, which greatly advanced grapevine research. This comprehensive assembly significantly contributes to our understanding of grape genetics and improves the precision of breeding efforts. However, most of the cultivated grapevines are highly heterozygous (Gaut et al., 2018; Xiao et al., 2023; Zhou et al., 2017; Zhou et al., 2019), which poses a challenge in studying their genomes. With the advent of Chromium sequencing technology and high-throughput chromatin conformation capture (Hi-C), an increasing number of whole genome sequences of plants have been determined, making the assembly and annotation of diploid genomes a routine task (Djari et al., 2024). For ancient grape germplasm resources, phylogenetic studies of Eurasian species of native grapes are even more challenging to carry out due to the lack of high-quality reference genome sequences and genome annotation information. In the current study, we constructed high-quality haplotype-resolved genomes of two representative ancient grape varieties, red and white Munage, providing important resources for comparative genomics of Eurasian grape species.

The Munage grape, an ancient variety extensively cultivated in Xinjiang (Fig. 2C), is renowned for its high-quality berries. These grapes known for their large size, thin skin, and plump flesh are highly valued as table grapes. Our assembled haplotypes were compared with the latest reference genome (PN_T2T) (Shi et al. 2023), and the findings provided clear evidence of high collinearity and significant differences among haplotypes (Fig. 1B). The estimated genome size of *V. vinifera* is approximately 500 Mb (Jaillon et al. 2007) and our findings revealed that the genome the genome size of WM and RM to be 537.74 Mb and 497 Mb, respectively in line with the recently published grape assemblies. The DNA heterozygosity level of both the genomes are 3.31%, indicating a high level of heterozygsity, like other grape varieties including ‘Chasselas’ and ‘Ugni Blanc’ (Djari et al. 2024). A high BUSCO score (> 98.3) in all haplotypes of Munage clearly indicated high-quality assembly and accurate annotation with low fragmentation and duplications (Wang et al. 2023). We annotated 33,942 (WM1), 34,034 (WM2), 35,292 (RM1), and 35,007 (RM2) genes, with TE annotations ranging between 67.15% and 66.56%. The annotated genes and TE annotations in previous grape assemblies, including ‘Shanputao’ (*V. amurensis*), ‘Noble’ (*M. rotundifolia*), ‘PN40024’. v2, and ‘Sultanina’ v2, were lower than what is in the current study, indicating the significance of our study in terms of assembly quality (Wang et al. 2023). The collinear analysis uncovered a range of structural variations, such as gap, inversions, duplications and translocations, in the genomes of the grape varieties and the reference genome.

Based on the population genetic analyses, we found that Munage has close relationship to Eurasia domesticated grapes. The single-branched structure of all cultivated grapes on the tree suggested that they may be of the same origin (Fig. 2A), while the PCA results showed that there were also significant differences between Munage and other cultivation differences under different artificial domestication selections (Fig. 2B). Based on our selection analysis, we found that significant differences in many regions across 19 chromosomes occurred during domestication process (Fig. 3A). Furthermore, genomic regions on Chr2 and Chr10 were identified by CLR analysis. During the domestication process, the selection process is largely driven by favoring traits that are desirable to humans (Lamoureux et al. 2006). This process results in the accumulation of genetic variants that enhance the desired characteristics. Over time, the genetic makeup of domesticated species diverges from their wild counterparts, reflecting the selective pressures exerted during domestication (Kui et al. 2020). In this study, the PBS analysis with multiple populations showed some selective sweep regions in the chromosomes. Significant differentiation across the chromosome region in the population provides strong evidence of natural selection over time in Munke grapes.

In recent years, research on grapes has received increasing attention as the cultivation and production of grapes in China has been rising exponentially. By using haplotype-resolved assemblies, we constructed haplotype genomes of Munage grapes to compare similarities and identify differences between RM and WM. These comparisons will help to understand the role of genomic regions which affect levels of gene expression, and key phenotypes, including berry skin color, one of the key appearance features determining the commercial value of grapevine. The recent genomic study of ‘Yan73’ provided insights into the color development mechanism in grapevines (Zhang *et al*., 2023). The white berries are assumed to have emerged from independent mutations (Kobayashi et al., 2004). It is assumed that RM originated by the bud mutation in WM. In our findings, we provided scientific evidence through IBD analysis. In our analysis of SNP variation followed by gene annotation of WM and RM, we found that *PMAT2* and *MYB123* on chromosome 12 and 13 (Fig. 4A) might be the possible genes which contribute to color in development of RM. Based on homology it was found that *PMAT2* is important for color development in *Nicotiana tabacum* (Fujiwara *et al*., 1997). The MYBs and UFGT were the genes identified as genes related to color development in ‘Yan73’ by genome assembly approach (Zhang *et al*., 2023). As a transcription factor family, MYB is identified as an important regulator of anthocyanins in plants, including grapes (Kobayashi *et al*., 2004; This et al., 2007). There is no clear evidence of the involvement of *MYB123* genes in grape skin color, whereas *AcMYB123* was identified as a prerequisite for anthocyanin production in kiwifruit (Wang et al., 2019a). Consistently, some homologues genes *TT2-like R2R3-MYBs* in other crops including *Fragaria ananassa* (Schaart et al., 2013) and *Prunus persica* (Uematsu et al., 2014) have been identified to play crucial roles in anthocyanin production.

## Materials and methods

### Plant Materials and sampling

An ancient grape variety with specific traits, a standard mother plant of red Munage over 30 years old, and a standard mother plant of white Munage grapes over 80 years old, were selected from various locations in Xinjiang, China. We extracted high-quality DNA from the leaves to generate the high-quality haplotype-resolved genome assemblies. To obtain this, we employed PacBio whole-genome high-fidelity (HiFi) 100× long-read-length sequencing, high-throughput chromosome conformation capture (Hi-C) proximity linkage technology, and Illumina short-read-length sequencing. We performed genome annotation using second-generation Illumina transcriptome sequencing data from different tissues of Munage. The details of the genetic resources used are presented in Table S3.

### DNA sequencing and Library preparation

The CTAB method was used to extract high-quality genomic DNA from leaf tissue samples. The concentration and purity of the DNA were assessed using a Qubit fluorometer from Thermo Fisher Scientific. Gel electrophoresis was performed to verify DNA integrity. To sequence the DNA of the standard mother plant of the Xinjiang native ancient grape variety Munage, third generation of PacBio Sequel CCS 100× with HiFi data volume of up to 50Gb and two cells was used. For PacBio HiFi sequencing, a standard SMRTbell library was constructed using the SMRTbell Express Template Prep Kit 2.0, following the manufacturer’s recommendations (Pacific Biosciences, CA, USA). Additionally, the Munage genome haplotypes were assembled using second-generation 50× Hi-C combination sequencing technology.

### Total RNA extraction, library preparation and sequencing

To perform RNA sequencing, we utilized the NEBNext® Ultra™ II Directional RNA Library Prep Kit for Illumina® (New England Biolabs, MA, USA) to extract total RNA from various plant parts, including the leaves, pulp, stalk, and root of the Munage grapevine. Subsequently, paired end reads of 150 base pairs were generated on the Illumina NovaSeq platform. The sample details are presented in Table S3.

### Genome comparison between haplotypes and reference gene

For genome comparison, we first aligned the WM1, WM2, RM1, RM2, and PN_T2T genomes using Minimap2 (Li, 2018) (v.2.21) and indexed the resulting BAM files with SAMtools (v1.4) (Danecek et al., 2021). Next, to identify synteny and structural rearrangements between the genomes, we used SyRI (v.1.7.0) (Goel et al., 2019). Finally, we visualized the alignment results using ‘plotsr’ (Goel and Schneeberger, 2022).

### SNP calling and filtering

We utilized resequencing data from a total of 59 grape samples, which included 10 wild grapes (*Vitis vinifera* subsp. *sylvestris*) from Europe (EU), 10 wild grapes from the Middle East (ME), 27 domesticated grapes (Other), three white Munage, and six red Munage. Additionally, three samples of Muscadine grapes were used as an outgroup. In this study, we generated resequencing data for nine Munage samples, while the remaining data was obtained from the National Center for Biotechnology Information (NCBI). To perform quality control on the resequencing data, we utilized fastp (v.0.21) with the default parameters (Chen, 2023). Subsequently, the filtered paired-end data was mapped to the PN_T2T complete reference genome, and SNPs were called from the resequencing data of 59 samples using GTX (v2.2.1) software for further analysis. To minimize false positives, we applied filtering to the data using VCFtools (AlbersCornelis et al., 2011). Specifically, we removed genotypes with a genotype quality less than 20 (option-minGQ 20), SNPs with more than two alleles (options -min-alleles 2 --max-alleles 2), and SNPs with more than 20% missing genotypes (option -max-missing 0.8). Indels were also filtered by VCFtools using command --remove-indels.

### Gene Ontology Enrichment Analysis

We downloaded the eukaryotic database provided by eggNOG for sequence alignment (Huerta-Cepas et al., 2019). HMMER3 was used to search each protein sequence in the eggNOG database. Each HMM match corresponded to an eggNOG OG with a functional annotation, providing preliminary annotation information. Subsequently, phmmer was used to search each protein sequence within the set of eggNOG proteins corresponding to the best matching HMM. The best match result for each sequence was then stored as a seed ortholog, which was used to obtain other orthologous genes. The final annotation result was the functional description of the orthologous genes that corresponded to the protein sequence used for the search. Finally, the obtained results were used for gene enrichment analysis by TBtools (Chen et al., 2020).

### Phylogeny and population structure

Principal component analysis was conducted using PLINK (Purcell et al., 2007) based on the obtained results. Phylogenetic trees were constructed using FastTree (v2.1.10) (Piñeiro et al., 2020) based on GTR models. Population structure was estimated using Admixture (v1.3.0) (Alexander et al., 2009) with K values ranging from 2 to 10, based on nuclear SNPs. IBD was calculated using PLINK with parameter: --genome.

### Selective sweep

To detect signatures of selective sweeps in Munage populations, we employed the Composite Likelihood Ratio (CLR) method implemented in SweeD (v3.2.1) (Pavlidis et al., 2013). The top 1% of likelihood values were used to identify potential selective sweeps within highly differentiated genomic regions. The first step involved partitioning the population and chromosomes. Next, we determined the length of each chromosome and set the size of the scanning window. Chromosome lengths were obtained from the VCF file header. To ensure uniform segmentation, we divided each chromosome into 10 kb grids, resulting in 100 grids per megabase. Finally, we ran SweeD and visualized the results after generating the output file. We conducted Population Branch Statistic (PBS) analysis using PBScan software (https://github.com/thamala/PBScan) (Hämälä and Savolainen, 2019) to detect evidence of selective sweeps among the closely related populations of Munage and others, with ME serving as the outgroup. We sorted the PBS output results in descending order and considered the top 1% as outliers for Gene Ontology (GO) enrichment analysis.

### Population genetic analyses

Sequence similarity (*D*_*xy*_) was calculated by applying the Python script popgenWindows.py (available at https://github.com/simonhmartin/genomics_general) to non-overlapping windows of 5 kb. To assess the absolute nucleotide diversity differences, comparisons were made between all pairs of groups (WM, n = 3; RM, n = 6).

### Transcriptome differential gene analysis

In this study, we used 3 WM leaves as a control, 3 WM peels, and 9 RM peels, and used WM2 as the reference genome for differential expression analysis of RNA-seq data. We used HISAT2’s ‘hisat2-build’ (v. 2.2.1) (Kim et al., 2019) to create an index for the haplotype of the reference genome. The transcriptome data were then aligned with the reference genome using HISAT2. SAMtools was employed to convert and sort the resulting SAM files into BAM files, and to create an index for the BAM files. Concurrently, we used ‘gffread’ (v.0.12.7) (Pertea and Pertea, 2020) to convert the gene annotation GFF3 file to GTF format. Next, we utilized the ‘featureCounts’ (Liao et al., 2014) software to accurately count reads by comparing the positions of each feature in the genome regions with the alignment positions of each base in the reads or fragments, thereby obtaining the gene expression value matrix result file.

Subsequently, we used DESeq2 (Love *et al*., 2014) to calculate the differential fold changes based on the gene expression value matrix results, obtaining the differential fold changes and significance *p*-values for each gene. Here, we aimed to filter the genes based on |log2FC| ≥ 2 and padj < 0.01, and to distinguish upregulated and downregulated genes using “up” and “down,” respectively.

## Author Contributions

Conceptualization: HX, XW,YZ, HZ, XS and FZ; Data curation: HZ, XS, FZ, YV, HX; Data analysis: HZ, XS, FZ, XW, SC, JQ,ZL,YL,MZ and TZ; Funding acquisition: XW and HZ; Methodology: HZ, XS, FZ, SZ,CZ,; Project administration: HZ; Resources: HZ, FZ, XZ and XW; Validation: HZ, XS, and FZ; Writing, review and editing, HZ, XS, YV and HX; Supervision: HX, XW, YZ,.

## Data availability

All raw data have been deposited in the NCBI Sequence Read Archive under project number PRJNA1158437 and the National Genomics Data Center (NGDC) Genome Sequence Archive (GSA) (https://ngdc.cncb.ac.cn/gsa/), with BioProject number PRJCA029988. The assembly and annotation have been deposited in zenodo: https://doi.org/10.5281/zenodo.13732355.

