## Supplemental Fig 1-4 for "Haplotype-resolved assemblies provide insights into genomic makeup of the oldest grapevine cultivar (Munage) in Xinjiang"

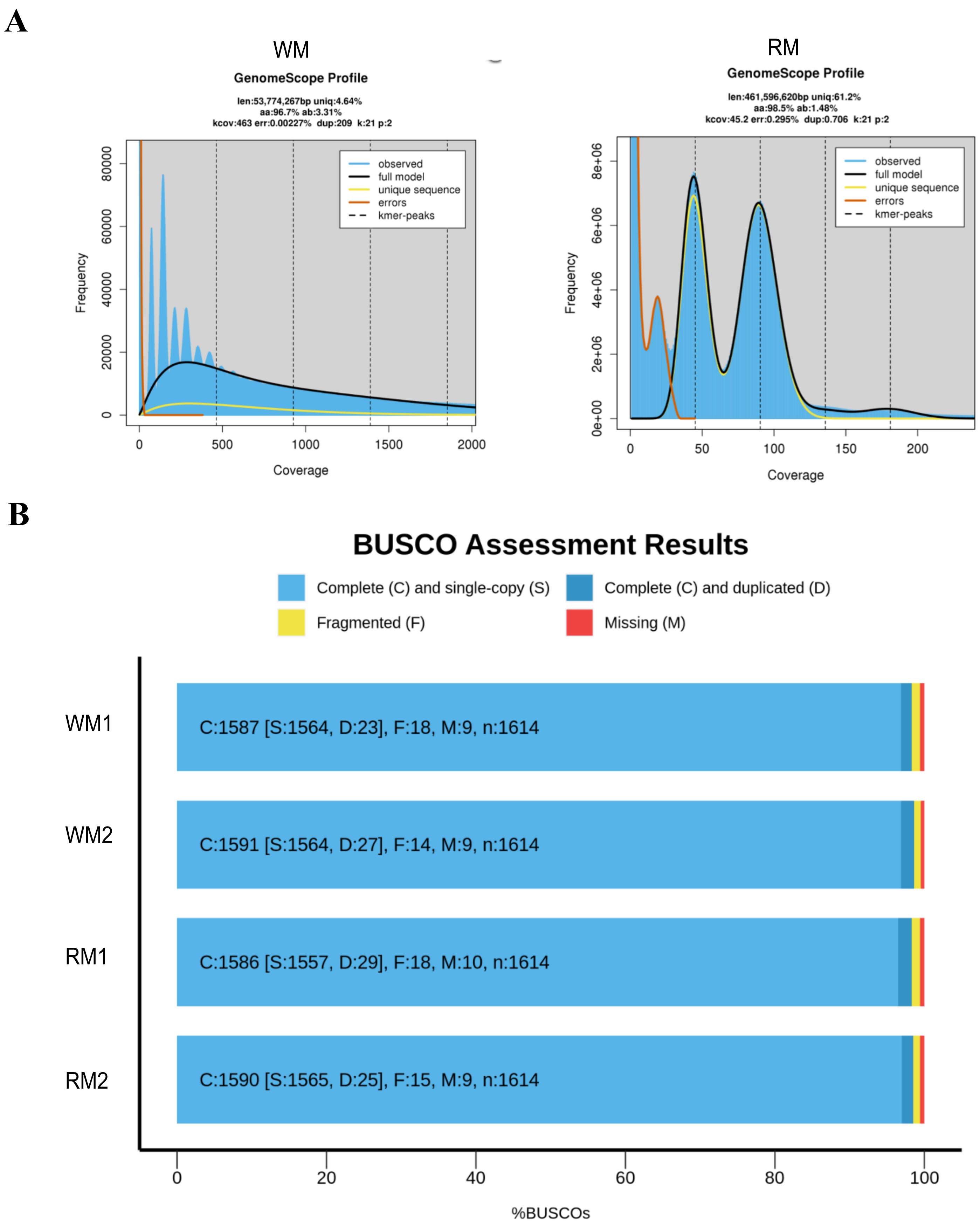


Figure S1 (A) Genome size and heterozygosity assessment using K-mer analysis.
(B) Genome completeness assessment using BUSCO.

A

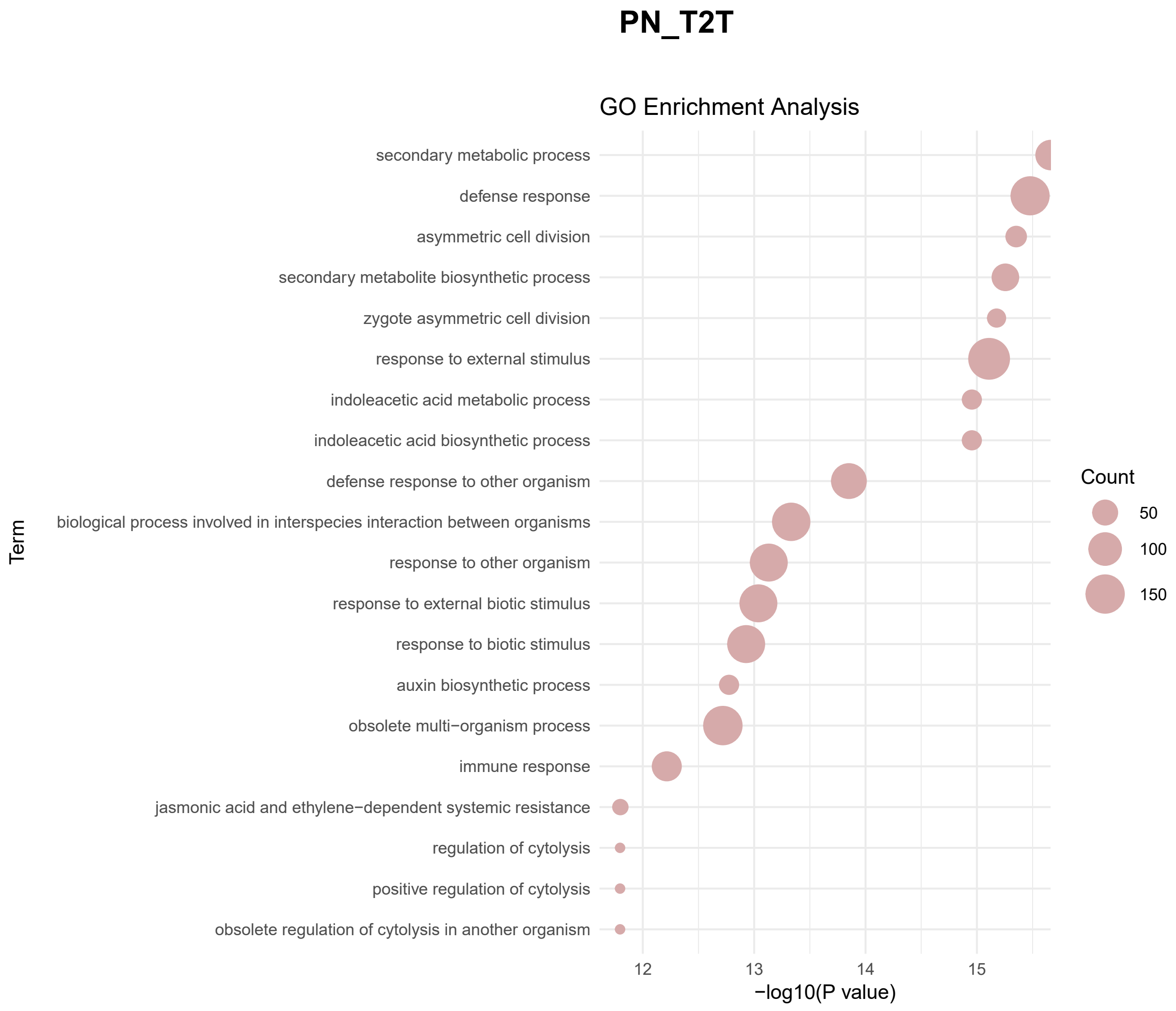
PN_T2T
GO Enrichment Analysis
secondary metabolic process
defense response
asymmetric cell division
secondary metabolite biosynthetic process
zygote asymmetric cell division
response to external stimulus
indoleacetic acid metabolic process
indoleacetic acid biosynthetic process
defense response to other organism
biological process involved in interspecies interaction between organisms
response to other organism
response to external biotic stimulus
response to biotic stimulus
auxin biosynthetic process
obsolete multi−organism process
immune response
jasmonic acid and ethylene−dependent systemic resistance
regulation of cytolysis
positive regulation of cytolysis
obsolete regulation of cytolysis in another organism
Term
12
13
14
15
−log10(P value)
Count
50
100
150

B

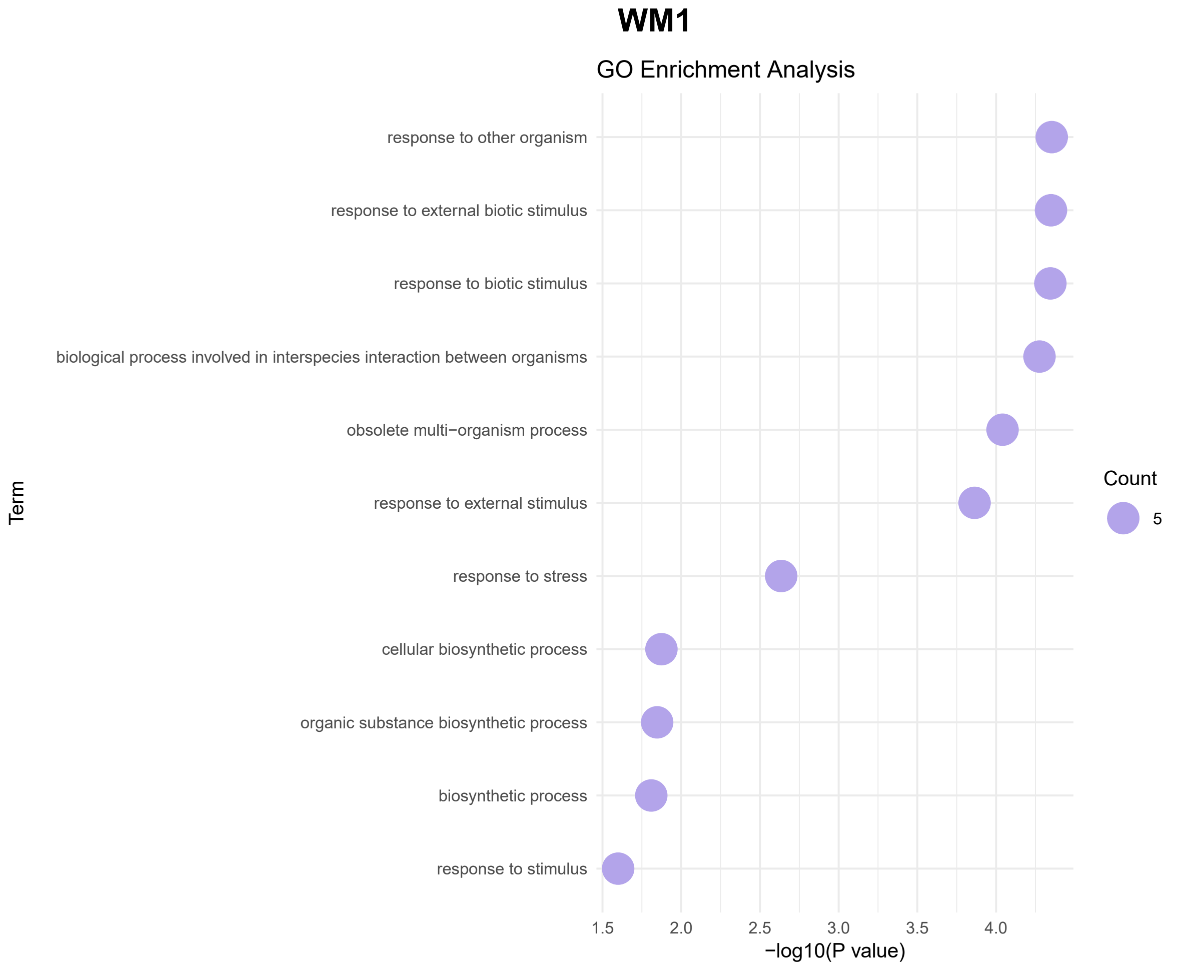
WM1
GO Enrichment Analysis
response to other organism
response to external biotic stimulus
response to biotic stimulus
biological process involved in interspecies interaction between organisms
obsolete multi−organism process
response to external stimulus
response to stress
cellular biosynthetic process
organic substance biosynthetic process
biosynthetic process
response to stimulus
Term
1.5
2.0
2.5
3.0
3.5
4.0
−log10(P value)
Count
5

## RM1

C

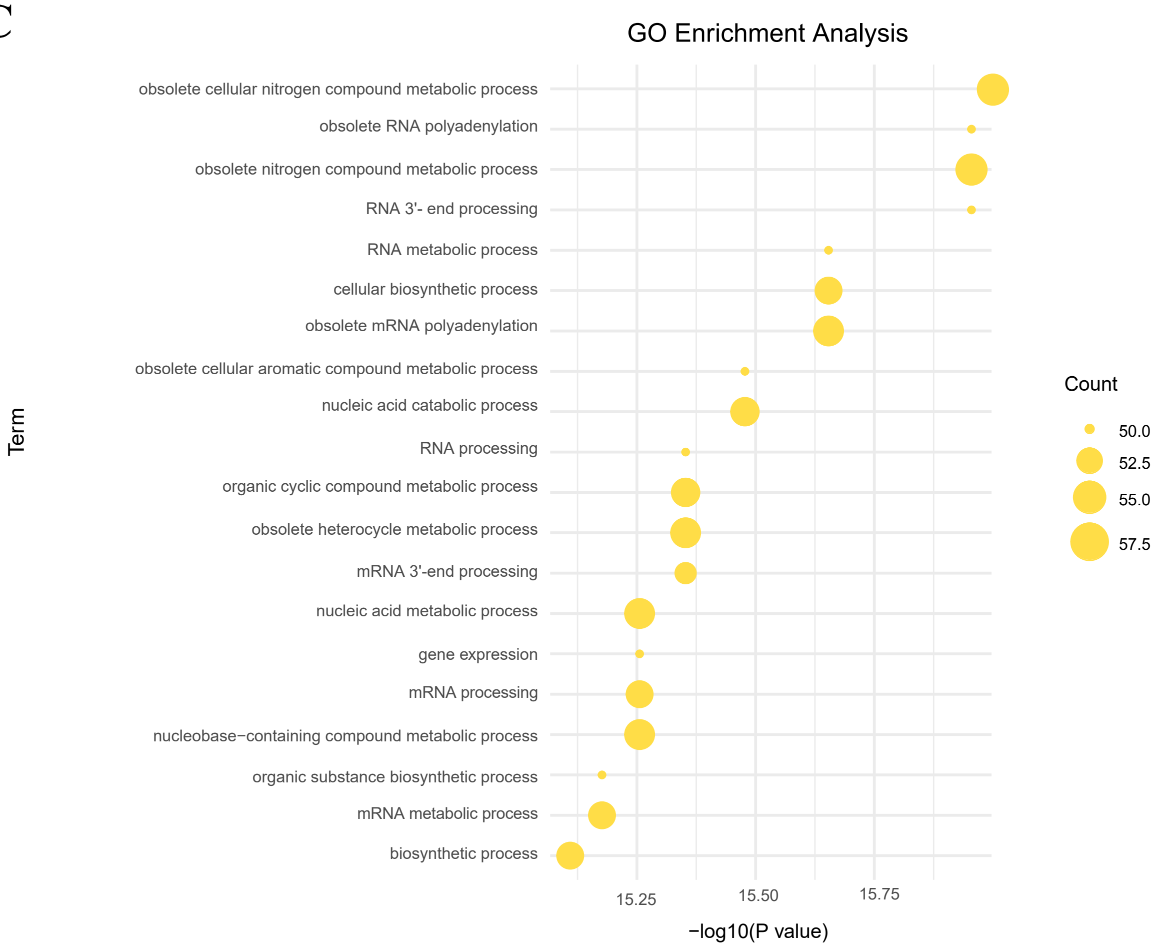


## RM2

D

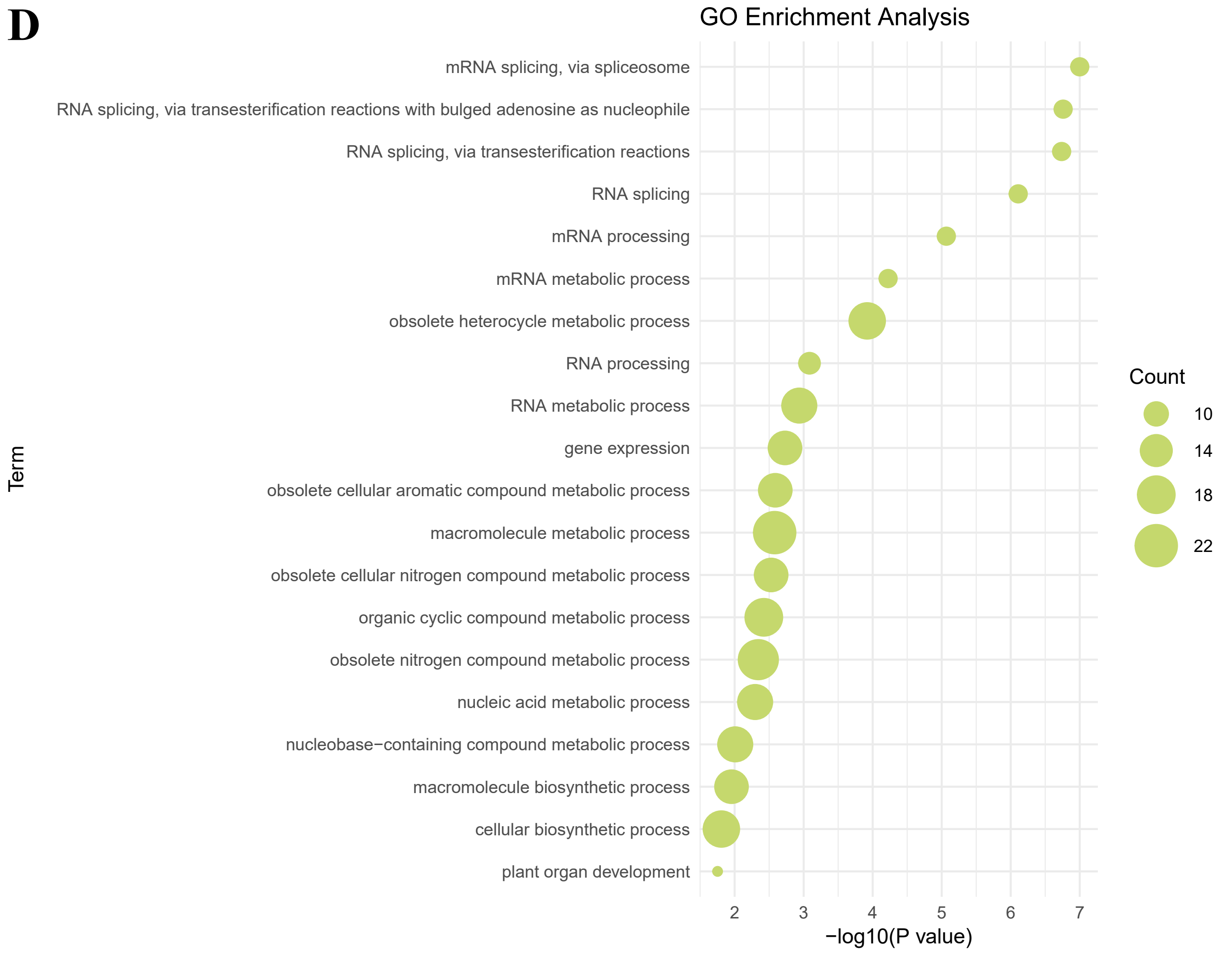


Figure S2 (A) (B) (C) (D) GO enrichment analysis of unique homologous genes in PN_T2T, WM1, RM1, and RM2.

A

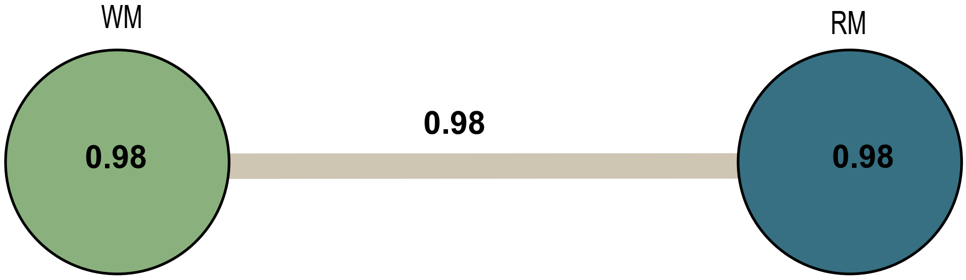


Figure S3 (A) IDB analysis of WM and RM. The values within the circles indicate the average IBD between individuals within each group, with larger circles reflecting a higher number of individuals. The values on the gray lines represent the average IBD between individuals in the two groups.

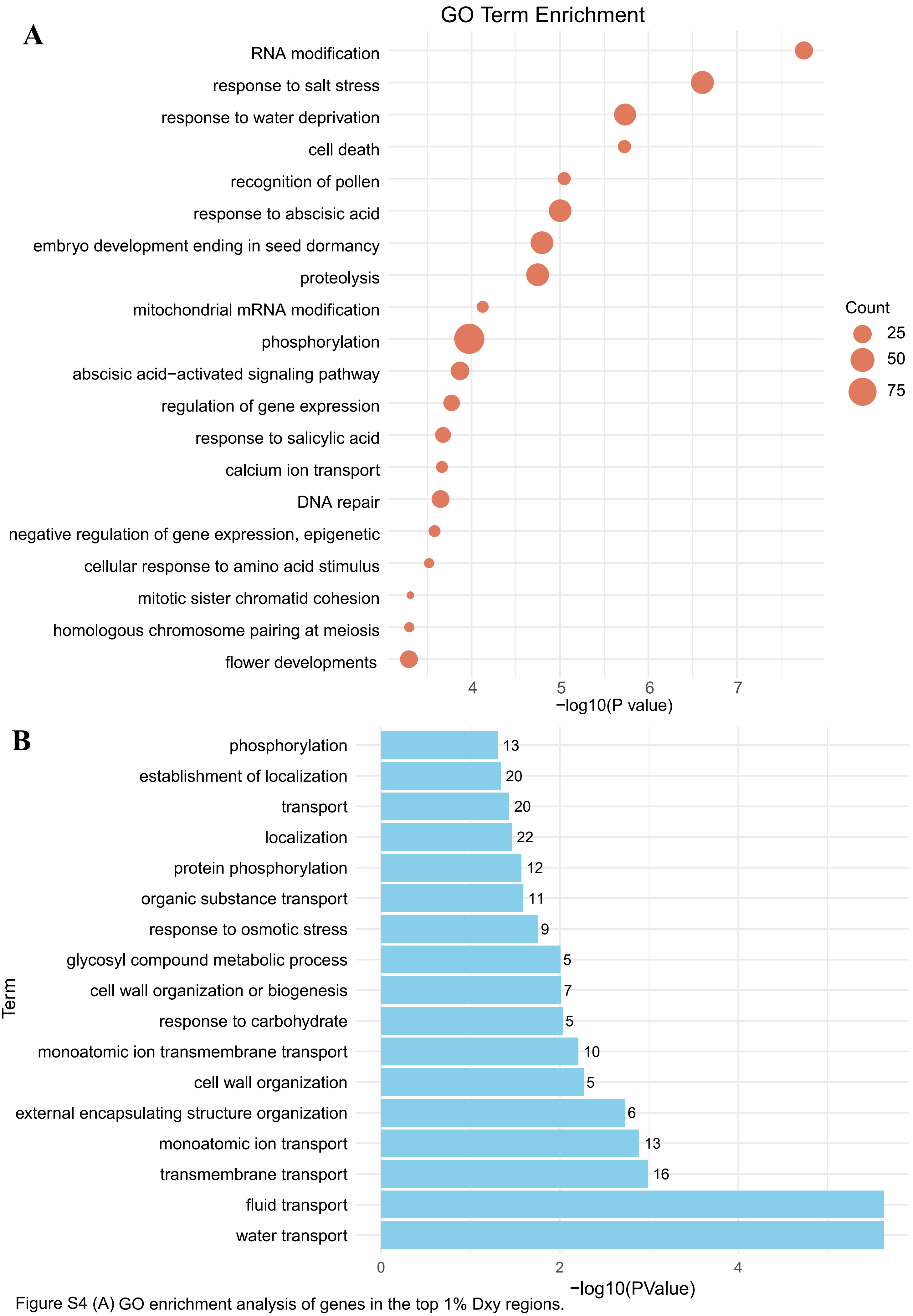


Figure S4 (A) GO enrichment analysis of genes in the top 1% Dxy regions.
(B) GO enrichment analysis of upregulated genes.
