## Supplemental Table 1-3 for "Haplotype-resolved assemblies provide insights into genomic makeup of the oldest grapevine cultivar (Munage) in Xinjiang"

**Table S1. Comparison of chromosome length WM1, WM2, RM1 and RM2 assembly.**

| Chr_ID | WM1_Length (bp) | WM2_Length(bp) | RM1_Length(bp) | RM2_Length(bp) |
|---|---|---|---|---|
| WM_hap1_chr1 | 24828091 | 24323354 | 24400844 | 24715939 |
| WM_hap1_chr2 | 21453684 | 21614472 | 21458218 | 21494222 |
| WM_hap1_chr3 | 21956811 | 21593469 | 21536987 | 22006528 |
| WM_hap1_chr4 | 27128911 | 26331393 | 26303377 | 27148028 |
| WM_hap1_chr5 | 27039085 | 27554711 | 26969450 | 27557495 |
| WM_hap1_chr6 | 25111505 | 24986057 | 25008310 | 25052721 |
| WM_hap1_chr7 | 31173380 | 30529185 | 31731389 | 30587366 |
| WM_hap1_chr8 | 24311910 | 25340795 | 25762510 | 24158373 |
| WM_hap1_chr9 | 23234080 | 26743558 | 24817437 | 23004635 |
| WM_hap1_chr10 | 26367376 | 27294045 | 25852394 | 26206440 |
| WM_hap1_chr11 | 22244874 | 19954744 | 20024093 | 20402234 |
| WM_hap1_chr12 | 23293471 | 25185014 | 24829081 | 23249738 |
| WM_hap1_chr13 | 29582808 | 28442042 | 29828677 | 28352613 |
| WM_hap1_chr14 | 29639168 | 28831715 | 29636154 | 28817980 |
| WM_hap1_chr15 | 22030944 | 22111255 | 21856749 | 22288342 |
| WM_hap1_chr16 | 24572247 | 23339235 | 23522504 | 24469924 |
| WM_hap1_chr17 | 20749501 | 20463297 | 20156923 | 19676922 |
| WM_hap1_chr18 | 37681528 | 37105468 | 37733526 | 37078949 |
| WM_hap1_chr19 | 25703744 | 27610568 | 28088058 | 24351175 |

**Table S2. Comparison of genomic features of WM1, WM2, RM1 and RM2 assemblies.**

| | WM1 | WM2 | RM1 | RM2 |
|---|---|---|---|---|
| Total sequence length (bp) | 488103118 | 489354377 | 489516681 | 480619624 |
| Number of chromosomes | 19 | 19 | 19 | 19 |
| Contig N50 (Mb) | 24.32 | 26.33 | 15.55 | 17.08 |
| Annotated centromere | 19 | 19 | 19 | 19 |
| Annotated telomere | 36 | 36 | 34 | 34 |
| The number of gene | 33942 | 34034 | 35292 | 35007 |
| Repeat content (%) | 67.04% | 67.15% | 67.02% | 66.56% |
| BUSCO | 98.3% | 98.6% | 98.3% | 98.5% |

**Table S3. Grape samples used in the analysis:** Samples numbered 1 to 50 are the whole genome sequencing data obtained in previous studies, and others are the data obtained in this study.

| | Name | Species | Group | Note |
|---|---|---|---|---|
| 1 | SRR22585228 | *Vitis californica* | OUT | WGS |
| 2 | SRR22585227 | *Vitis californica* | OUT | WGS |
| 3 | SRR22585216 | *Vitis californica* | OUT | WGS |
| 4 | ERR6359144 | *V. vinifera* subsp.*sylvestris* | EU | WGS |
| 5 | ERR6359145 | *V. vinifera* subsp. *sylvestris* | EU | WGS |
| 6 | ERR6359493 | *V. vinifera* subsp. *sylvestris* | EU | WGS |
| 7 | ERR6359504 | *V. vinifera* subsp. *sylvestris* | EU | WGS |
| 8 | ERR6359506 | *V. vinifera* subsp. *sylvestris* | EU | WGS |
| 9 | SRR5891608 | *V. vinifera* subsp. *sylvestris* | EU | WGS |
| 10 | SRR5891613 | *V. vinifera* subsp. *sylvestris* | EU | WGS |
| 11 | SRR5891768 | *V. vinifera* subsp. *sylvestris* | EU | WGS |
| 12 | SRR5891764 | *V. vinifera* subsp. *sylvestris* | EU | WGS |
| 13 | SRR22585225 | *V. vinifera* subsp. *sylvestris* | EU | WGS |
| 14 | SRR22585211 | *V. vinifera* subsp. *sylvestris* | ME | WGS |
| 15 | SRR22585210 | *V. vinifera* subsp. *sylvestris* | ME | WGS |
| 16 | SRR22585207 | *V. vinifera* subsp. *sylvestris* | ME | WGS |
| 17 | SRR22585206 | *V. vinifera* subsp. *sylvestris* | ME | WGS |
| 18 | SRR22585198 | *V. vinifera* subsp. *sylvestris* | ME | WGS |

| | | | | |
|---|---|---|---|---|
| 19 | SRR22585203 | *V. vinifera* subsp. *sylvestris* | ME | WGS |
| 20 | SRR22585202 | *V. vinifera* subsp. *sylvestris* | ME | WGS |
| 21 | SRR22585214 | *V. vinifera* subsp. *sylvestris* | ME | WGS |
| 22 | SRR22585218 | *V. vinifera* subsp. *sylvestris* | ME | WGS |
| 23 | SRR22585217 | *V. vinifera* subsp. *sylvestris* | ME | WGS |
| 24 | CRR493691 | *V. vinifera* subsp. *vinifera* | Other | WGS |
| 25 | CRR493822 | *V. vinifera* subsp. *vinifera* | Other | WGS |
| 26 | CRR493839 | *V. vinifera* subsp. *vinifera* | Other | WGS |
| 27 | CRR493967 | *V. vinifera* subsp. *vinifera* | Other | WGS |
| 28 | CRR494957 | *V. vinifera* subsp. *vinifera* | Other | WGS |
| 29 | CRR495198 | *V. vinifera* subsp. *vinifera* | Other | WGS |
| 30 | CRR495592 | *V. vinifera* subsp. *vinifera* | Other | WGS |
| 31 | CRR495848 | *V. vinifera* subsp. *vinifera* | Other | WGS |
| 32 | CRR496293 | *V. vinifera* subsp. *vinifera* | Other | WGS |
| 33 | CRR496605 | *V. vinifera* subsp. *vinifera* | Other | WGS |
| 34 | CRR496710 | *V. vinifera* subsp. *vinifera* | Other | WGS |
| 35 | CRR497182 | *V. vinifera* subsp. *vinifera* | Other | WGS |
| 36 | CRR498694 | *V. vinifera* subsp. *vinifera* | Other | WGS |
| 37 | CRR498888 | *V. vinifera* subsp. *vinifera* | Other | WGS |
| 38 | ERR8014963 | *V. vinifera* subsp. *vinifera* | Other | WGS |

| | | | | |
|---|---|---|---|---|
| 39 | ERR8014964 | *V. vinifera* subsp. *vinifera* | Other | WGS |
| 40 | SRR8835168 | *V. vinifera* subsp. *vinifera* | Other | WGS |
| 41 | ERR3046968 | *V. vinifera* subsp. *vinifera* | Other | WGS |
| 42 | VV_70 | *V. vinifera* subsp. *vinifera* | Other | WGS |
| 43 | VV_97 | *V. vinifera* subsp. *vinifera* | Other | WGS |
| 44 | VV_100 | *V. vinifera* subsp. *vinifera* | Other | WGS |
| 45 | VV_102 | *V. vinifera* subsp. *vinifera* | Other | WGS |
| 46 | VV_147 | *V. vinifera* subsp. *vinifera* | Other | WGS |
| 47 | VV_167 | *V. vinifera* subsp. *vinifera* | Other | WGS |
| 48 | VV_178 | *V. vinifera* subsp. *vinifera* | Other | WGS |
| 49 | VV_198 | *V. vinifera* subsp. *vinifera* | Other | WGS |
| 50 | VV_225 | *V. vinifera* subsp. *vinifera* | Other | WGS |
| 51 | GB1 | *V. vinifera* subsp. *vinifera* | WM | WGS |
| 52 | GB4 | *V. vinifera* subsp. *vinifera* | WM | WGS |
| 53 | GB9 | *V. vinifera* subsp. *vinifera* | WM | WGS |
| 54 | GB2 | *V. vinifera* subsp. *vinifera* | RM | WGS |
| 55 | GB3 | *V. vinifera* subsp. *vinifera* | RM | WGS |
| 56 | GB5 | *V. vinifera* subsp. *vinifera* | RM | WGS |
| 57 | GB7 | *V. vinifera* subsp. *vinifera* | RM | WGS |
| 58 | GB8 | *V. vinifera* subsp. *vinifera* | RM | WGS |

| | | | | |
|---|---|---|---|---|
| 59 | GB10 | *V. vinifera* subsp. *vinifera* | RM | WGS |
| 60 | WM1 | *V. vinifera* subsp. *vinifera* | WM | HIFI |
| 61 | RM1 | *V. vinifera* subsp. *vinifera* | RM | HIFI |
| 62 | WM1 | *V. vinifera* subsp. *vinifera* | WM | HIC |
| 63 | RM1 | *V. vinifera* subsp. *vinifera* | RM | HIC |
| 64 | BMY4 | *V. vinifera* subsp. *vinifera* | WM | RNA-seq (leaf) |
| 65 | BMY5 | *V. vinifera* subsp. *vinifera* | WM | RNA-seq (leaf) |
| 66 | BMY6 | *V. vinifera* subsp. *vinifera* | WM | RNA-seq (leaf) |
| 67 | BMP4 | *V. vinifera* subsp. *vinifera* | WM | RNA-seq (pericarp) |
| 68 | BMP5 | *V. vinifera* subsp. *vinifera* | WM | RNA-seq (pericarp) |
| 69 | BMP7 | *V. vinifera* subsp. *vinifera* | WM | RNA-seq (pericarp) |
| 70 | HMP4 | *V. vinifera* subsp. *vinifera* | RM | RNA-seq (pericarp) |
| 71 | HMP5 | *V. vinifera* subsp. *vinifera* | RM | RNA-seq (pericarp) |
| 72 | HMP6 | *V. vinifera* subsp. *vinifera* | RM | RNA-seq (pericarp) |
| 73 | HMBP4 | *V. vinifera* subsp. *vinifera* | RM | RNA-seq (pericarp) |

| | | | | |
|---|---|---|---|---|
| 74 | HMBP5 | *V. vinifera* subsp. *vinifera* | RM | RNA-seq (pericarp) |
| 75 | HMBP6 | *V. vinifera* subsp. *vinifera* | RM | RNA-seq (pericarp) |
| 76 | HMHP4 | *V. vinifera* subsp. *vinifera* | RM | RNA-seq (pericarp) |
| 77 | HMHP5 | *V. vinifera* subsp. *vinifera* | RM | RNA-seq (pericarp) |
| 78 | HMHP6 | *V. vinifera* subsp. *vinifera* | RM | RNA-seq (pericarp) |
